# Graph-Based Reconstruction and Analysis of Disease Transmission Networks using Viral Genomic Data

**DOI:** 10.1101/2022.07.28.501873

**Authors:** Ziqi Ke, Haris Vikalo

## Abstract

Understanding the patterns of viral disease transmissions helps establish public health policies and aids in controlling and ending a disease outbreak. Classical methods for studying disease transmission dynamics that rely on epidemiological data, such as times of sample collection and duration of exposure intervals, struggle to provide desired insight due to limited informativeness of such data. A more precise characterization of disease transmissions may be acquired from sequencing data that reveals genetic distance between viral populations in patient samples. Indeed, genetic distance between viral strains present in hosts contains valuable information about transmission history, thus motivating the design of methods that rely on genomic data to reconstruct a directed disease transmission network, detect transmission clusters, and identify significant network nodes (e.g., super-spreaders). In this paper, we present a novel end-to-end framework for the analysis of viral transmissions utilizing viral genomic (sequencing) data. The proposed framework groups infected hosts into transmission clusters based on reconstructed viral quasispecies; the genetic distance between a pair of hosts is calculated using Earth Mover’s Distance, and further used to infer transmission direction between the hosts. To quantify the significance of a host in the transmission network, the importance score is calculated by a graph convolutional auto-encoder. The viral transmission network is represented by a directed minimum spanning tree utilizing the Edmond’s algorithm modified to incorporate constraints on the importance scores of the hosts. Results on realistic synthetic as well as experimental data demonstrate that the proposed framework outperforms state-of-the-art techniques for the analysis of viral transmission dynamics.

**CCS CONCEPTS:**

- Applied computing → Bioinformatics.

**ACM Reference Format:** Ziqi Ke and Haris Vikalo. 2022. Graph-Based Reconstruction and Analysis of Disease Transmission Networks using Viral Genomic Data. In *Proceedings of The Seventh International Workshop on Computational Network Biology (CNB-MAC 2022)*. ACM, New York, NY, USA, 10 pages. https://doi.org/XXXXXXX.XXXXXXX

## 1 INTRODUCTION

Understanding the spread of a pathogen across a network of hosts assists in the development of effective public health interventions and disease prevention, containment and eradication strategies. Examples include studies of infectious disease transmissions in support of reconstructing a transmission network, detecting transmission clusters, and identifying super-spreaders in the network. Traditional methods for infectious disease outbreak analysis that rely on epidemiological data such as the time of testing and duration of exposure suffer from labour-intensive contact tracing [19], and generally struggle to provide desired insight due to limited informativeness of such data. For example, the time of testing is an unreliable indicator of the time of infection, especially for a disease that may be asymptomatic long after the infection (e.g., in the case of the original COVID-19 strain, the symptoms occur 2-14 days following the infection). With the advance of next-generation sequencing (NGS) technologies, rapid and accurate reconstruction of viral populations is feasible and affordable. Since genomic evolutionary distance between viral strains present in different hosts contains valuable information about transmission history, analysis of genomic data collected by NGS technologies may provide significant insight into disease transmission patterns.

Existing methods for studying disease transmission patterns can be classified as (i) epidemiological data driven, (ii) genomic data driven, and (iii) methods that rely on the combination of both epidemiological and genomic data. Prior work on utilizing epidemiological data includes a social network analysis of Mycobacterium Tuberculosis transmissions using patient medical records and contact interview forms [8]; stochastic mathematical models describing disease transmission process using both behavioral and environmental data [18]; analysis of social networks built utilizing existing clinical informatics resources aiming to explore the implications of patient-healthcare worker interactions on disease transmission [1]; study of a human contact network formed using close proximity interaction data, meant to provide insight into transmissions of an influenza-like disease during a typical day at an American high school [36]; a stochastic eco-epidemiological model for the analysis of dengue transmission using seasonal and spatial dynamics [33]; a 2009 H1N1 pandemic influenza transmission model estimated via Markov chain Monte Carlo sampling of demographic and clinical data [7]; likelihood-based methods for the analysis of influenza A(H1N1) transmission in a school population using clinical symptom and contact data [22]; mathematical modeling of the transmission of lumpy skin disease virus using direct and indirect contact information of cattle exhibiting typical clinical signs of the disease [27]; network models of transmission dynamics in wild animal and livestock populations using contact data [10]; a statistical inference method for the construction of the influenza A virus transmission tree in a college-based population utilizing epidemiological, clinical and contact tracing data [49]; a model enabling reconstruction of the full-spectrum dynamics of COVID-19 using epidemiological data [20]; statistical modeling for the reconstruction of transmission pairs for COVID-19 utilizing detailed demographic characteristics, travel history, social relationships and epidemiological timelines [46]; and a visualization technique based on individual reports of epidemiological data to construct disease transmission graphs for the COVID-19 epidemic [26].

As an alternative, methods that aim to go beyond traditional techniques for outbreak analysis by relying on genomic data have recently started gaining attention, including a graph based technique for reconstructing transmission trees utilizing genomic data from the early stages of the A/H1N1 influenza pandemic [24]; minimum spanning tree based methods for estimating relationships among individual strains or isolates in molecular epidemiology [38]; a Bayesian approach for reconstructing densely sampled outbreaks from whole-genome sequence data and inferring a transmission network via a Monte Carlo Markov chain [12]; an approach for analyzing genetic distance between pathogen strains to estimate routes of transmission in bacterial disease outbreaks [43]; a method for molecular detection of hepatitis C virus transmissions in outbreak settings [6]; stochastic epidemic models for investigating person-to-person communicable disease transmission with densely sampled genomic data [45]; a statistical framework to infer host-to-host transmissions built around a computationally efficient model of pathogen evolution [11]; algorithms to infer genetic relatedness, detect possible transmissions, and analyze clusters’ structure validated using real sequencing data from HCV outbreaks [16]; an approach incorporating shared genetic variants and phylogenetic distance data to identify transmission routes from pathogen deep-sequence data [44]; a graph-based method for modeling viral evolution and epidemic spread via evolutionary analysis of intra-host viral populations [40]; a Bayesian approach for transmission inference that explicitly models evolution of pathogen populations in an outbreak [28]; a statistical learning approach with a pseudo-evolutionary model to infer epidemiological links from deep sequencing data for human, animal and plant diseases [3], and an algorithm based on Earth mover’s distance for viral outbreak investigations utilizing raw NGS reads [30].

Methods relying on the combination of epidemiological and genomic data include a minimum spanning tree model to identify the history of transmission of hepatitis C virus in an outbreak [41]; a maximum-likelihood approach for the analysis of HIV-1 transmissions utilizing clinical, epidemiological and phylogenomic data [15]; a stochastic infectious disease model for the analysis of the spread of infectious salmon anaemia among salmon farms utilizing genetic and space-time data [4]; a likelihood-based framework drawing upon temporal, geographical and genomic data in an epidemic of avian influenza A (H7N7) in The Netherlands in 2003 [47]; Bayesian inference frameworks to reconstruct most likely transmission patterns and infection dates for the analysis of UK epidemics of foot-and-mouth disease virus [9, 25, 32]; a statistical framework to infer key epidemiological and mutational parameters by simultaneously estimating the phylogenetic and transmission tree in an outbreak of foot-and-mouth disease [48]; a statistical method exploiting both pathogen sequences and collection dates to reveal dynamics of densely sampled outbreaks for the analysis of the 2003 Singaporean outbreak of Severe Acute Respiratory Syndrome (SARS) [23]; a Bayesian inference method for the reconstruction of HIV transmission trees from viral sequences and uncertain infection time data [31]; a systematic Bayesian transmission network model to reconstruct the transmission network of the foot-and-mouth disease epidemic in Japan in 2010 [21]; a Bayesian methodology that uses contact data for the inference of transmission trees in a statistically rigorous manner, alongside genomic data and temporal data [5], and an analysis of mumps virus transmission in the US utilizing epidemiological data from public health investigations and mumps virus whole genome sequences [42].

In this paper, we present an end-to-end framework for the reconstruction and interpretation of disease transmissions from genomic data. The proposed framework aims to first divide patients into different clusters, where patients insider the same cluster are infected by the same type of viral variant of the infectious disease; to this end, we utilize a modified version of the viral quasispecies reconstruction method TenSQR [2]. Next, the genetic distance between a pair of patients inside the same cluster is calculated based on the Earth Mover’s Distance between corresponding *k*-mer distributions [30, 34], which reflects the minimal amount of work that must be performed to transform one distribution into another by moving ‘distribution mass’ around. Third, possible transmission directions between pairs of patients are determined using the calculated genetic distances. Fourth, the importance score of every patient is calculated via a custom-designed graph convolutional auto-encoder inspired by [37]. Auto-encoders are neural networks that can be trained to automatically extract salient low-dimensional representations of high-dimensional data in an unsupervised manner [17]; graph convolutional auto-encoders are a variant of autoencoders specifically designed to analyze graph-structured data. Finally, a directed minimum spanning tree, incorporating the local and global information provided by the genetic distances and the importance scores, is constructed by imposing importance scores constraints in the classical Edmonds’ algorithm [13]. The final version of the paper will provide link to a Github repository containing the codes.

Our main contributions are summarized as follows:

- We developed an end-to-end computational framework that utilizes sequencing data to detect disease transmission clusters, reconstruct a directed disease transmission network, and quantify the significance of a viral host within the transmission network.
- Our proposed method reconstructs the directed disease transmission network by leveraging not only the local information captured by the genetic distances but also the global information provided by the importance scores of each host extracted by a designed graph convolutional auto-encoder.
- We conducted extensive experiments on experimental COVID-19 data and semi-experimental hepatitis C virus data, and compared the results with existing state-of-the-art methods, demonstrating the ability of the proposed framework to efficiently and accurately reconstruct a disease transmission network, outperforming state-of-the-art methods.

## 2 METHODS

### 2.1 Problem Formulation

The aims of the framework for disease transmission network analysis developed in this paper include:

1. Discovery of transmission clusters, where the clusters collect hosts infected by the same pathogen variant.
2. Inference of a directed transmission network of hosts based on genomic information about viral pathogens infecting them.
3. Identification of super-spreaders/critical hosts in the network.

To detect clusters of hosts/patients, we adapt to the current problem the viral quasispecies reconstruction method TenSQR we previously co-developed in [2]. Specifically, we first successively cluster hosts into communities represented by the consensus sequences of cluster-specific viral genomes, and then construct a weighted directed acyclic graph *G* = (*V*, *E*, **w**) for each cluster, where *V* is the set of nodes representing hosts, *E* is the set of edges indicating disease transmission directions, and **w** is the set of edge weights. For instance, node *v_i_* denotes the *i^th^* host, edge *e_ij_* = (*v_i_*, *v_j_*) indicates that the *j^th^* host may have been infected by the *i^th^* host, and weight *w_ij_* is the genetic distance between the pathogens infecting the two hosts. Following [30], we represent the genetic distance separating two pathogens via the Earth Mover’s Distance between corresponding *k*-mer distributions. While the genetic distances provide useful local information about transmissions, we further design a graph convolutional auto-encoder to obtain global information in form of importance scores quantifying how influential hosts are in the network. The importance scores are in turn used to guide the search for a directed minimum spanning tree revealing transmissions along the network. Fig. 1 illustrates the architecture of the proposed end-to-end framework.

**Figure 1:**
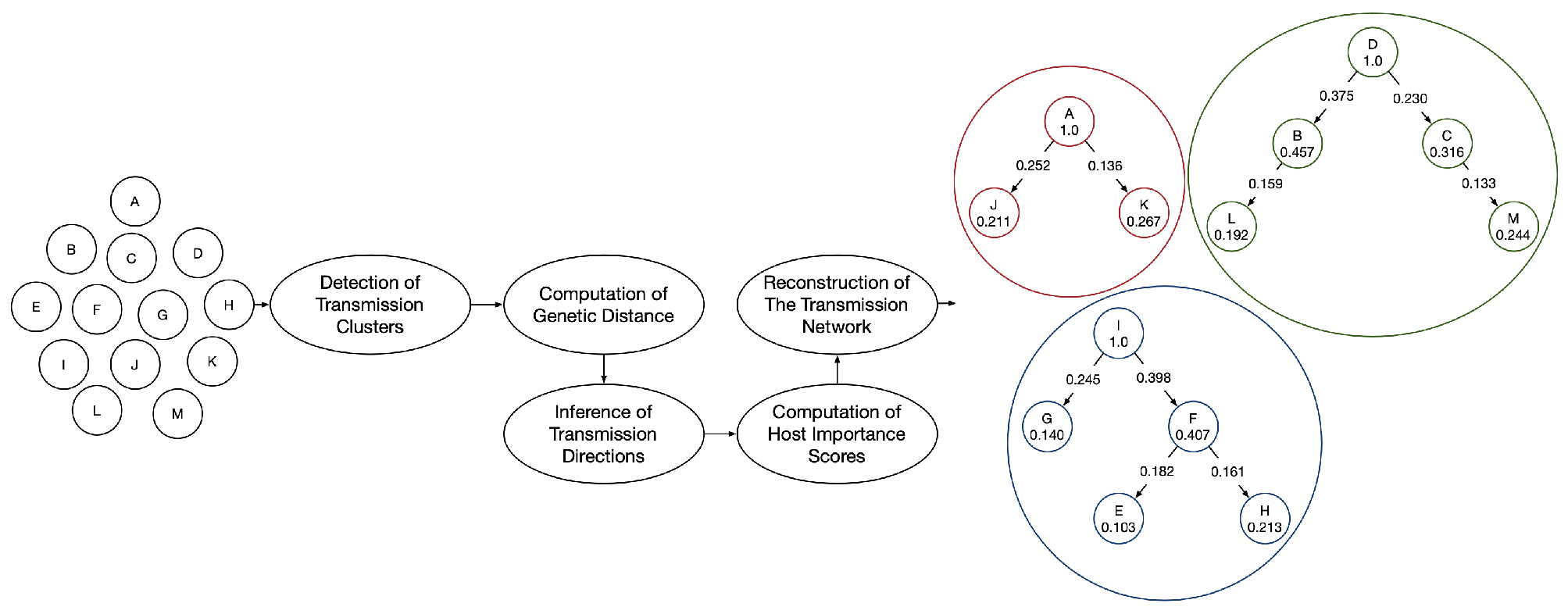
Architecture of the proposed end-to-end framework for disease transmission analysis. Differently colored nodes illustrate three disease transmission clusters, with arrows showing the transmission directions. The numbers shown in nodes and on edges are the importance scores and the genetic distances, respectively.

### 2.2 Detection of Transmission Clusters

As already mentioned, the method used to detect disease transmission clusters builds upon TenSQR proposed in our prior work [2]. Let *P* denote an *n* × *l* pathogen matrix where *n* is the number of hosts and *l* is the (maximum) length of the viral genomes; note that the entries in *P* may be erroneous due to sequencing errors. After organizing hosts into transmission clusters, where hosts in the same cluster are infected by the same variant type of the pathogen, the consensus sequence is formed to represent each cluster. The reconstructed consensus sequences form a *k* × *l* consensus matrix *C*, where *k* denotes the number of transmission clusters. To ensure the distances between nucleotides are consistent, we denote nucleotides by 4-dimensional standard unit vectors 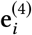, 1 ≤ *i* ≤ 4, with 0s in all positions except the *i^th^* one that has value 1 (e.g., 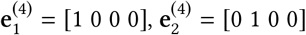, and so on). The pathogen matrix *P* can be re-written as a binary tensor 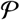 whose fibers represent nucleotides and horizontal slices correspond to pathogen strains. The transmission cluster detection can then be formulated as a tensor factorization problem since 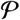 can be thought of as being obtained by multiplying an *n* × *k* pathogen strain membership indicator matrix *M* and a binary tensor 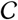 that encodes the consensus sequence of each cluster. Fibers of 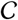 are standard unit vectors 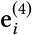 representing alleles, while each lateral slice of 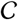 is one of the *k* consensus sequences representing cluster centroids. Note that the pathogen strain membership indicator matrix *M* has standard unit vectors 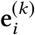, 1 ≤ *i* ≤ *k*, for rows; if the *j^th^* row of **M** is 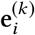, that indicates the *j^th^* host is assigned to the *i^th^* transmission cluster. To proceed, we formulate the transmission network clustering problem as a collection of *k* −1 tensor factorization problems; after each factorization, pathogens associated with the most dominant transmission cluster are removed from 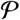 and the factorization (of smaller dimension) is performed anew until only one cluster remains.

Fig. 2 illustrates the detection of transmission clusters. To formalize the tensor factorization representation of the problem, let **P** ∈ {0,1}^n×41^ and **C** ∈ {0,1}^*k*×*l*^ denote the mode-1 unfoldings of tensors 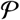 and 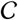, respectively. The transmission network clustering problem can be written as

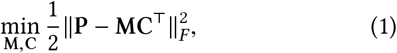

where ║ · ║_F_ denotes the Frobenius norm. This is a non-convex optimization that can be approximately solved via alternating minimization. For further details regarding finding a solution to the above optimization problem and thus successively identifying pathogens associated with the transmission clusters, we refer a reader to [2].

**Figure 2:**
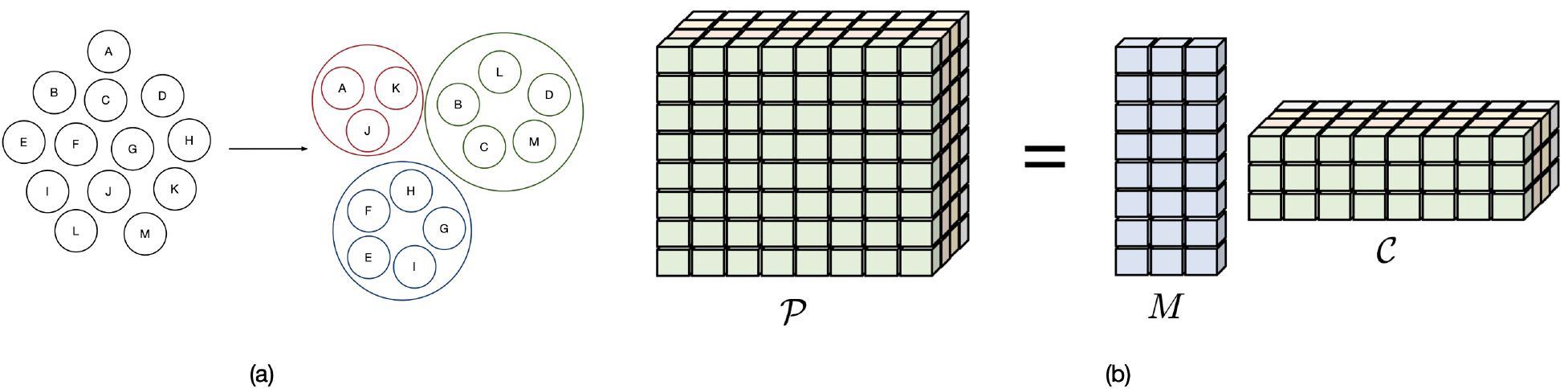
(a) Detecting transmission clusters. (b) Identifying viral species via tensor factorization.

### 2.3 Computation of Genetic Distances

Given a cluster of hosts, we build a graph in which the edge weights reflect genetic similarities between pathogens infecting the hosts. The Earth Mover’s Distance (EMD) used to measures the difference between two probability distributions is calculated as a solution to the transportation problem (i.e., the Monge-Kantorovich problem), which is readily formalized as a linear program [34, 35]. Let

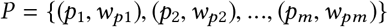

denote the normalized *k*-mer frequencies of a viral strain, where *p_i_* represents the *i^th^ k*-mer and *w_pi_* is its frequency. Similarly, let us denote the normalized *k*-mer frequencies of another viral strain by

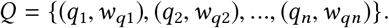

Let *D* = [*d_ij_*] be the shortest path distance between *k*-mers *p_i_* and *p_j_* in the undirected De Bruijn graph formed by all the pathogens in a transmission cluster. [Fig. 3 illustrates a sample de Bruijn graph constructed from two sequences with *k* = 4.] The Earth Mover’s Distance between *P* and *Q* is defined as

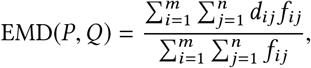

where the total flow 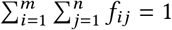. The aforementioned linear program is focused on finding *F* = [*f_ij_*] minimizing

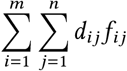

subject to

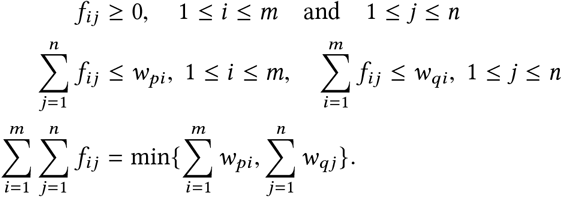

where *f_ij_* denotes the flow between *p_i_* and *q_j_*.

**Figure 3:**
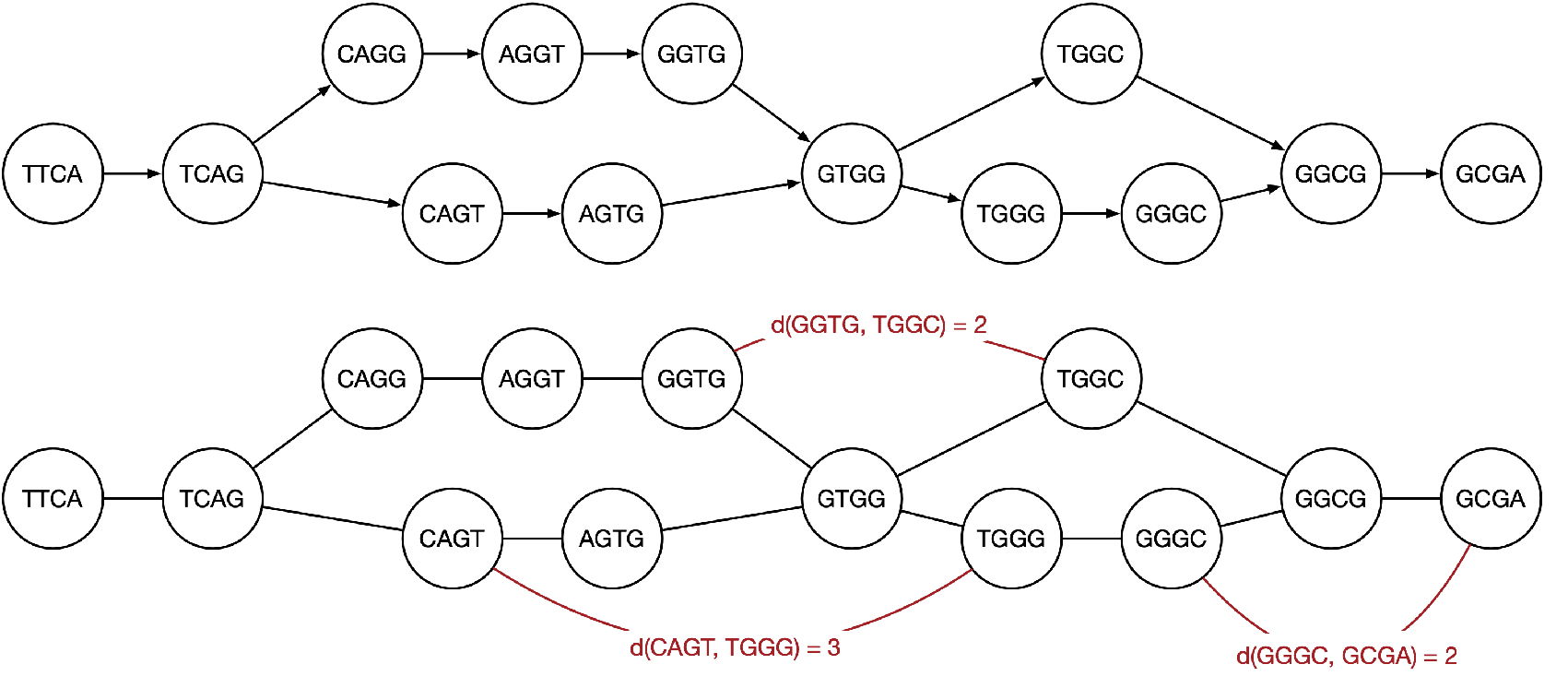
An example of a de Bruijin graph constructed from sequences TTCAGTGGGCGA and TTCAGGTGGCGA. A few selected pairwise distances between *k*-mers are also indicated.

### 2.4 Inference of Transmission Directions

For two hosts *A* and *B* with normalized *k*-mer frequencies 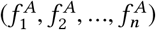 and 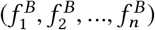, where *n* denotes the total number of unique *k*-mers in *A* and *B*, the maximum mean *k*-mer distribution is defined as [30]

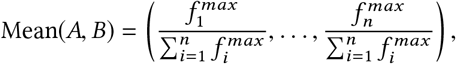

where 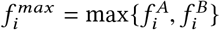 for all *i* = 1, 2,…, *n*. Then the transmission direction between hosts A and B is assumed to be from *A* to *B* if

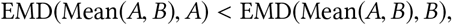

and from B to A otherwise; as before, EMD(·, ·) denotes the Earth Mover’s Distance between its arguments. Fig. 4 illustrates inference of the transmission direction between hosts *A* and *B*.

**Figure 4:**
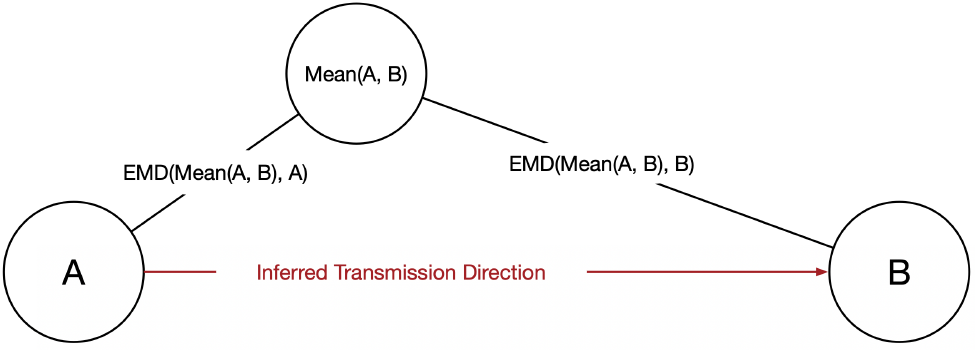
An illustration of inferring transmission direction between host A and host B.

### 2.5 Computing Host Importance Scores via Graph Convolutional Auto-Encoder

Consider a host network represented by the weighted directed graph *G* = (*V*, *E*, **w**), where |*V*| = *n* is the number of hosts and where weights **w** reflect genetic distances between hosts. Let *X* denote the *n* × *n* matrix of distances between pathogens associated with the hosts, and let *A* denote the adjacency matrix of *G*. Motivated by [37], we design the graph convolutional encoder (GCE) that learns to construct an *n* × (*d* + 1) node embedding matrix |*Z*, *M*|, where | ***·*** | denotes matrix concatenation and the dimensions of *Z* and *M* are *n* × *d* and *n* × 1, respectively. The first *d* ≪ *f* dimensions of the embedding correspond to the latent feature representation of a host, whereas the last dimension corresponds to a mass parameter *m_i_* ∈ *R*^+^ of the host; note that *m_i_* is reflective of the impact of host *i* on the graph flow and thus signifies its importance in the spread of the disease. By drawing parallels to the Newton’s theory of universal gravitation [37], acceleration 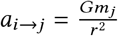 (where 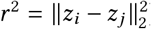) can be interpreted as an indicator of the likelihood that the pathogen associated with host *i* is a genetic ancestor of the pathogen associated with host *j*. A graph convolutional encoder with *L* layers, where *L* ≥ 2, and |*Z*, *M*| = *H*^(*L*)^ can be summarized as

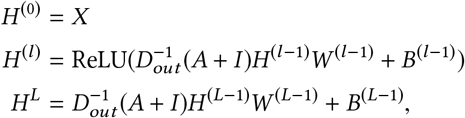

where *l* ∈ {1, 2,…, *L* – 1}, *D_out_* denotes the diagonal out-degree matrix of *A* + *I, I* represents the identity matrix, and *W* and *B* are learnable weight and bias matrix, respectively. In summary, a graph convolutional encoder learns node embeddings from *A* and *X* as |*Z*, *M*| = GCE(*A*, *X*). A graph decoder is then leveraged to reconstruct *A* from *Z* and *M* while allowing for asymmetric connectivity between hosts. The reconstructed adjacency matrix can be represented as 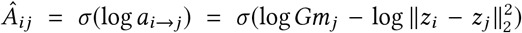, where *σ*(*x*) = 1/(1 + *e*^-*x*^) denotes the sigmoid function and *m_j_* is the importance score of the *j^th^* host; note that, in general, *Â_ij_* ≠ *Â_ji_*. The loss function in the form of weighted cross entropy is given by *L* = −∑_*i,j*_ *A_ij_* log *Â_ij_*. We build a two-layer graph convolutional encoder with layer dimensions set to 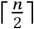 and 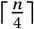. The learning rate is 0.001, and the number of training epoch is 200.

### 2.6 Reconstructing the Transmission Network

In the final stage of our framework, the weighted directed graph is pruned to obtain the disease transmission network. To this end, we modify the classical Edmond’s algorithm [13] to enable leveraging both the local and global information about the transmission dynamics; the former is provided by the genetic distances while the latter is given by the node importance scores. In particular, given a weighted directed graph *G* = (*V*, *E*, **w**, **s**), where *V* is the set of vertices representing the hosts, *E* is the set of directed edges, **w** is the set of edge weights, and **s** is the set of importance scores of the hosts, we would like to find a directed minimum spanning tree *T* having the smallest weight and satisfying the constraint that transmission directions are from the hosts with higher importance scores to the hosts with lower importance scores. Let *w*(*e*) be the weight on edge *e*, *w*(*u, v*) denote the weight on the edge from *u* to *v*, and *s*(*u*) be the importance score of host *u*. A pseudo-code of our modification of the Edmond’s algorithm, which for convenience we refer to as Edmond+Score, is given below.

#### Algorithm [Edmond+Score]

1. Select a host with the highest importance score as the root *r*, and remove all edges whose destination is *r*.
2. For each node *v* except *r*, keep the edge with the lowest weight incoming to *v* from *π*(*v*), where *s*(*π*(*v*)) ≥ *s*(*v*), and remove other edges whose destination is *v*. If the set of edges *P* = {(*π*(*v*), *v*)|*v* ∈ *V* \ *r*} does not contain cycles, the desired directed minimum spanning tree is found. Otherwise, go to step (3).
3. If there is at least one cycle in *P*, select one such cycle and denote it by *C*. Define a new weighted directed graph *G*′ = (*V*′, *E*′, **w**′, **s**) and treat *C* as a single (virtual) node, *v*_*C*_. For any edge from *u* ∉ *C* to *v* ∉ *C*, add a new edge *e* = (*u*, *v*_*C*_) to *E*′ and add *w*′(*e*) = *w*(*u*, *v*) – *w*(*π*(*v*), *v*) to **w**′, where *π*(*v*) is the source of *v* in *C*. For any edge from *u* ∈ *C* to *v* ∉ *C*, add a new edge *e* = (*v_C_*, *v*) to *E*′ and add *w*′(*e*) = *w*(*u*, *v*) to **w**′. For an edge from *u* ∉ *C* to *v* ∉ *C*, add a new edge *e* = (*u, v*) to *E*′ and add *w*′(*e*) = *w*(*u, v*) to **w**’.
4. For all edges incoming to *v_C_*, identify (*u*, *v*) with the lowest edge weight and remove (*π*(*v*), *v*) from *C* to break the cycle.
5. Repeat steps (3) and (4) until all the cycles in *G* are broken.

Fig. 5 shows an example of pruning a weighted directed graph with the Edmond+Score algorithm.

**Figure 5:**
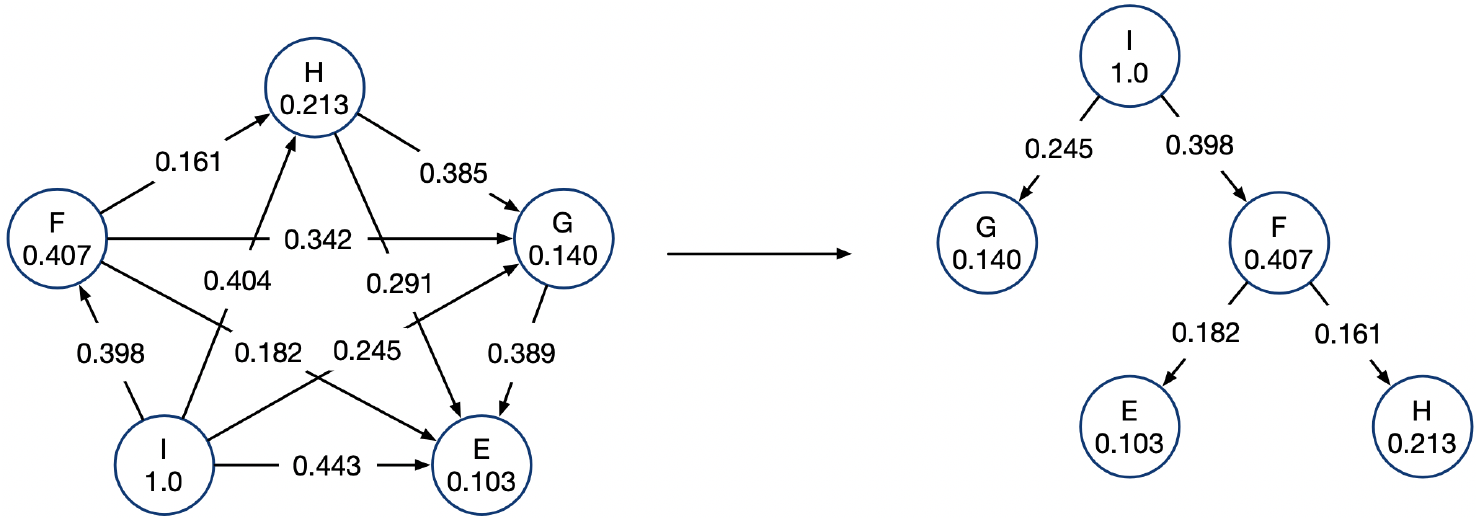
An example of a directed minimum spanning tree obtained by leveraging both local and global information about the transmission dynamics.

## 3 RESULTS

In this section, we compare the performance of the introduced methodology with state-of-the-art techniques for transmission network inference; for convenience, we refer to the proposed end-to-end framework as AutoNet, as it relies on scores provided by an auto-encoder to infer the network.

### 3.1 Performance on Semi-Experimental Hepatitis C Virus Data

Performance of AutoNet is first tested on semi-experimental hepatitis C virus data and compared with state-of-the-art methods including k-MED [30], QUENTIN [40] and MinDist [6]. The semi-experimental data is generated by judiciously introducing mutations in the experimentally obtained sequencing data of pathogens collected from a cardiac surgeon and five patients infected by the surgeon [14]. In particular, pathogens associated with the surgeon and five patients are represented by a sequence of 188 nucleotides encompassing the first hypervariable region (HVR-1) at the junction between the coding regions for envelope glycoproteins E1 and E2. The mean and standard deviation of the Hamming distance between the 188-nucleotide sequences are 11.0 and 3.84, respectively. We follow the steps below to repeatedly synthesize disease transmission networks:

1. Randomly select two out of the five patients infected by the surgeon.
2. For each selected patient, generate a new pair of (fictitious) patients by randomly inducing 11 mutations into the respective 188-nucleotides-long pathogen sequences.
3. Randomly select one patient from each pair generated in step (2), and repeat step (2) for the selected two patients. This creates a tree-like (depth 3) disease transmission network that we aim to reconstruct using only the 188-nucleotides-long pathogen sequence information.

The considered methods are compared in terms of the transmission direction accuracy, defined as the ratio of the number of pairs of hosts with correctly predicted transmission direction to the total number of host pairs, and the source identification accuracy, defined as the fraction of the transmission networks with correctly predicted sources. Table 1 shows the results on a semi-experimental dataset containing 2000 disease transmission networks. As can be seen, AutoNet achieves both the highest transmission direction accuracy as well as the highest source identification accuracy.

**Table 1:**
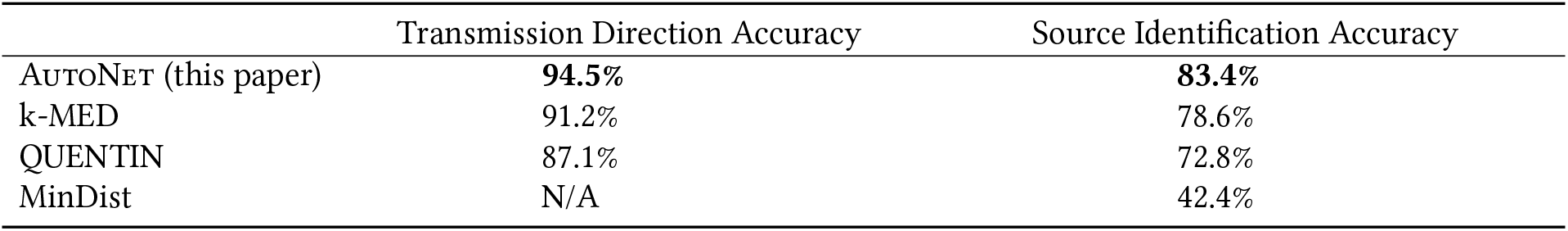
The performance comparison of AutoNet and the competing methods on the semi-experimental dataset with 2000 disease transmission networks. Note that MinDist cannot infer transmission directions (hence, N/A).

### 3.2 Performance on Experimental City-Level COVID-19 Data

Next, we compare performance of AutoNet with the selected state-of-the-art methods on experimental city-level COVID-19 data from the Global Initiative on Sharing All Influenza Data (GISAID) [39]. The GISAID promotes rapid sharing of data from all influenza viruses and the coronavirus causing COVID-19. This includes genetic sequences and related clinical and epidemiological data associated with human viruses, as well as geographical and species-specific data associated with the avian and other animal viruses, to help researchers understand how viruses evolve and spread during epidemics and pandemics (https://www.gisaid.org). For each host/patient, the experimental COVID-19 data includes the collection date, submission date, location, gender, patient age, virus variant type, specimen source, sequencing technology, assembly method, sequencing coverage, originating lab, submitting lab and so on. We focus on data collected in London between January 1st and January 31st of 2021, which includes 4252 hosts and 2 virus variant types, B.1.1.7 and B.1.177. We first test the performance of TenSQR [2] adapted to the task of of detecting transmission clusters using the reported virus variant types as the ground truth, and then report the spanning trees reconstructed by AutoNet and k-MED [39]. Since the sequences of reported pathogens associated with the 4252 hosts are shorter than the sequence lengths of the COVID-19 reference genome, the host sequences are first aligned to the COVID-19 reference genome (NCBI Reference Sequence: NC_045512.2) using BLAST [29]; TenSQR is then deployed to detect transmission clusters. The accuracy, precision and recall of the clustering results achieved by TenSQR are 0.998, 0.995 and 0.984, respectively. The F-1 score and the area under the ROC curve (AUC) are 0.989 and 0.992, respectively. Figures 6 (a) and (b) illustrate the sub-graphs of the disease transmission networks reconstructed by AutoNet and k-MED, respectively. Note that we only show the results achieved by AutoNet and k-MED because QUENTIN could not complete the task in 48 hours, while MinDist cannot infer transmission directions.

**Figure 6:**
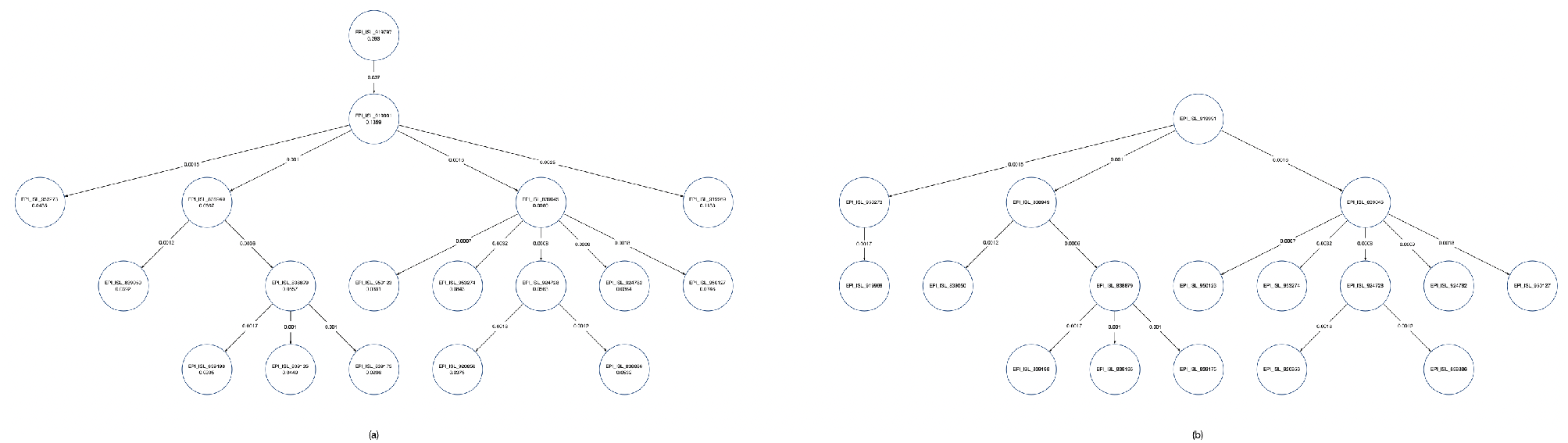
Experiments on city-level COVID-19 data. (a) The disease transmission network reconstructed by AutoNet; note the GISAID accession IDs and the importance scores for each host node. (b) The result of k-MED.

### 3.3 Performance on Experimental Country-Level COVID-19 Data

Finally, we compare performance of AutoNet and the considered state-of-the-art methods on experimental country-level COVID-19 data aiming to discover how the pathogens are transmitted between different cities. In addition to London’s, COVID-19 data collected in Alderley Edge, Milton Keynes, Cambridge, Glasgow, Oxford and Edinburgh between January 1st and January 31st of 2021 are also analyzed. The number of hosts present in the data for these additional cities is 16433, 9977, 7494, 4644, 1085 and 154, respectively. We rely on the consensus sequence of the dominant viral variant type to represent each city, and then perform the reconstruction task. The host sequences are aligned to the COVID-19 reference genome (NCBI Reference Sequence: NC_045512.2) before obtaining the consensus sequence for each city. Fig. 7 (a), (b) and (c) show the reconstruction results obtained by AutoNet, k-MED and QUENTIN, respectively. It is worth pointing out that our AutoNet and QUENTIN reconstruct similar networks, and that all the methods identify Oxford as the source of the spread.

**Figure 7:**
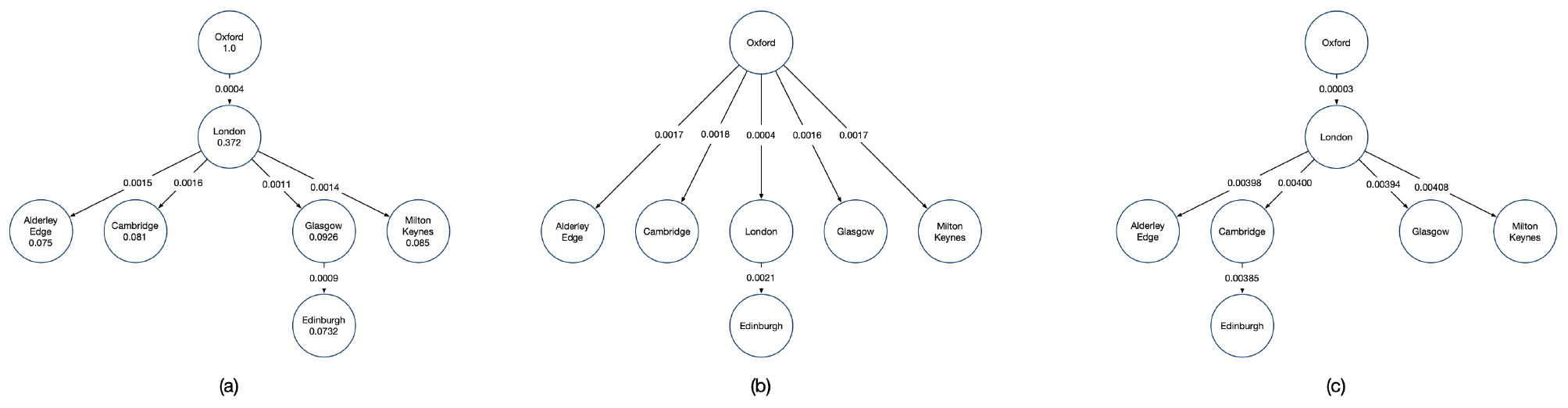
Experiments on country-level COVID-19 data. (a) The disease transmission network reconstructed by AutoNet. (b) The result of k-MED. (c) The result of QUENTIN.

## 4 CONCLUSIONS

In this paper we presented AutoNet, an end-to-end framework for the detection of disease transmission clusters, reconstruction of directed transmission networks, and the discovery of super-spreaders from genomic data. The framework first clusters infected viral hosts into groups where the hosts in a group are infected by the same pathogen variant. After quantifying similarity between pairs of viral hosts in a group, directions of transmission between hosts are estimated and the importance score of each host is calculated. Finally, a directed minimum spanning tree is reconstructed by leveraging both the local and global information about transmissions, provided by the genetic similarity between hosts and the hosts’ importance scores calculated via a graph autoencoder. Benchmarking on semi-experimental and real data shows that the proposed framework is able to reconstruct disease transmission networks efficiently and accurately, outperforming state-of-the-art competing techniques. Future work includes extending the proposed framework to enable joint processing of both genomic and epidemiological data.

## REFERENCES

[1] Gundlapalli Adi, Ma Xiulian, Benuzillo Jose, Pettey Warren, Greenberg Richard, Hales Joseph, Leecaster Molly, and Samore Matthew. 2009. Social network analyses of patient-healthcare worker interactions: implications for disease transmission. AMIA Annu Symp Proc (2009), 213/7.

[2] Soyeon Ahn, Ziqi Ke, and Haris Vikalo. 2018. Viral quasispecies reconstruction via tensor factorization with successive read removal. Bioinformatics 34, 13 (2018), i23–i31.

[3] M. Alamil, J. Hughes, K. Berthier, C. Desbiez, G. Thébaud, and S. Soubeyrand. 2019. Inferring epidemiological links from deep sequencing data: a statistical learning approach for human, animal and plant diseases. Phil. Trans.R. Soc. 374, 1775 (2019), 20180258.

[4] M Aldrin, T M Lyngstad, A B Kristoffersen, B Storvik, Ø Borgan, and P A Jansen. 2011. Modelling the spread of infectious salmon anaemia among salmon farms based on seaway distances between farms and genetic relationships between infectious salmon anaemia virus isolates. Journal of the Royal Society, Interface 8, 62 (09 2011), 1346–1356. https://doi.org/10.1098/rsif.2010.0737

[5] Finlay Campbell, Anne Cori, Neil Ferguson, and Thibaut Jombart. 2019. Bayesian inference of transmission chains using timing of symptoms, pathogen genomes and contact data. PLoS Comput Biol 15, 3 (2019), e1006930.

[6] David S. Campo, Guo-Liang Xia, Zoya Dimitrova, Yulin Lin, Joseph C. Forbi, Lilia Ganova-Raeva, Lili Punkova, Sumathi Ramachandran, Hong Thai, Pavel Skums, Seth Sims, Inna Rytsareva, Gilberto Vaughan, Ha-Jung Roh, Michael A. Purdy, Amanda Sue, and Yury Khudyakov. 2016. Accurate Genetic Detection of Hepatitis C Virus Transmissions in Out break Settings. The Journal of Infectious Diseases 213, 6 (5/15/2021 2016), 957–965. https://doi.org/10.1093/infdis/jiv542

[7] Simon Cauchemez, Achuyt Bhattarai, Tiffany L Marchbanks, Ryan P Fagan, Stephen Ostroff, Neil M Ferguson, and David Swerdlow. 2011. Role of social networks in shaping disease transmission during a community outbreak of 2009 H1N1 pandemic influenza. Proc Natl Acad Sci 108, 7 (2011), 2825–30.

[8] Victoria J. Cook, Sumi J. Sun, Jane Tapia, Stephen Q. Muth, D. Fermín Argüello, Bryan L. Lewis, Richard B. Rothenberg, and Peter D. McElroy. 2007. Transmission Network Analysis in Tuberculosis Contact Investigations. The Journal of Infectious Diseases 196, 10 (2007), 1517–1527.

[9] Eleanor M Cottam, Thébaud Gaël, Wadsworth Jemma, Gloster John, Mansley Leonard, Paton David J, King Donald P, and Haydon Daniel T. 2008. Integrating genetic and epidemiological data to determine transmission pathways of foot-and-mouth disease virus. Proc. R. Soc. B. 275, 1637 (2008), 887–895.

[10] Meggan E. Craft. 2015. Infectious disease transmission and contact networks in wildlife and livestock. Phil. Trans. R. Soc (2015), 370.

[11] Wilson DJ De Maio N, Wu C-H. 2016. SCOTTI: Efficient Reconstruction of Transmission within Outbreaks with the Structured Coalescent. PLoS Comput Biol 12, 9 (2016), e1005130.

[12] Xavier Didelot, Jennifer Gardy, and Caroline Colijn. 2014. Bayesian Inference of Infectious Disease Transmission from Whole-Genome Sequence Data. Molecular Biology and Evolution 31, 7 (5/15/2021 2014), 1869–1879. https://doi.org/10.1093/molbev/msu121

[13] J. Edmond. 1967. Optimum Branchings. Journal of Research of the National Bureau of Standards Section B 71B, 4 (1967), 233–240.

[14] JI Esteban, J Gómez, M Martell, B Cabot, J Quer, J Camps, A González, T Otero, A Moya, R Esteban, and et al. 1996. Transmission of hepatitis C virus by a cardiac surgeon. N Engl J Med 334, 9 (1996), 555–60.

[15] Martin Fisher, David Pao, Alison E Brown, Darshan Sudarshi, O Noel Gill, Patricia Cane, Andrew J Buckton, John V Parry, Anne M Johnson, Caroline Sabin, and Deenan Pillay. 2010. Determinants of HIV-1 transmission in men who have sex with men: a combined clinical, epidemiological and phylogenetic approach. AIDS 24, 11 (2010). https://journals.lww.com/aidsonline/Fulltext/2010/07170/Determinants_of_HIV_1_transmission_in_men_who_have.15.aspx

[16] Olga Glebova, Sergey Knyazev, Andrew Melnyk, Alexander Artyomenko, Yury Khudyakov, Alex Zelikovsky, and Pavel Skums. 2017. Inference of genetic relatedness between viral quasispecies from sequencing data. BMC genomics 18, Suppl 10 (12 2017), 918–918. https://doi.org/10.1186/s12864-017-4274-5

[17] Ian Goodfellow, Yoshua Bengio, and Aaron Courville. 2016. Deep Learning. MIT Press.

[18] Nicholas C. Grassly and Christophe Fraser. 2008. Mathematical models of infectious disease transmission. Nature Reviews Microbiology 6, 6 (2008), 477–487.

[19] MD Hall, M E J Woolhouse, and A Rambaut. 2016. Using genomics data to reconstruct transmission trees during disease outbreaks. Revue scientifique et technique (International Office of Epizootics) 35, 1 (2016), 287–296. https://doi.org/10.20506/rst.35.1.2433

[20] Xingjie Hao, Shanshan Cheng, Degang Wu, Tangchun Wu, Xihong Lin, and Chaolong Wang. 2020. Reconstruction of the full transmission dynamics of COVID-19 in Wuhan. Nature 584, 7821 (2020), 420–424. https://doi.org/10.1038/s41586-020-2554-8

[21] Yoko Hayama, Simson M Firestone, Mark A Stevenson, Takehisa Yamamoto, Tatsuya Nishi, Yumiko Shimizu, and Toshiyuki Tsutsui. 2019. Reconstructing a transmission network and identifying risk factors of secondary transmissions in the 2010 foot-and-mouth disease outbreak in Japan. Transbound Emerg Dis 66, 5 (2019), 2074–2086.

[22] Niel Hens, Laurence Calatayud, Satu Kurkela, Teele Tamme, and Jacco Wallinga. 2012. Robust Reconstruction and Analysis of Outbreak Data: Influenza A(H1N1)v Transmission in a School-based Population. American Journal of Epidemiology 176, 3 (2012), 196–203.

[23] Thibaut Jombart, Anne Cori, Xavier Didelot, Simon Cauchemez, Christophe Fraser, and Neil Ferguson. 2014. Bayesian Reconstruction of Disease Outbreaks by Combining Epidemiologic and Genomic Data. PLoS Comput Biol 10, 1 (2014), e1003457.

[24] T. Jombart, R M Eggo, P J Dodd, and F. Balloux. 2011. Reconstructing disease outbreaks from genetic data: a graph approach. Heredity 106, 2 (2011), 383–390. https://doi.org/10.1038/hdy.2010.78

[25] Max S. Y. Lau, Glenn Marion, George Streftaris, and Gavin Gibson. 2015. A Systematic Bayesian Integration of Epidemiological and Genetic Data. PLoS Comput Biol 11, 11 (2015), e1004633.

[26] Caiying Luo, Yue Ma, Pei Jiang, Tao Zhang, and Fei Yin. 2021. The construction and visualization of the transmission networks for COVID-19: A potential solution for contact tracing and assessments of epidemics. Scientific Reports 11, 1 (2021), 8605. https://doi.org/10.1038/s41598-021-87802-x

[27] Reuma Magori-Cohen, Yoram Louzoun, Yael Herziger, Eldad Oron, Alon Arazi, Eeva Tuppurainen, Nahum Y. Shpigel, and Eyal Klement. 2012. Mathematical modelling and evaluation of the different routes of transmission of lumpy skin disease virus. Veterinary Research 43, 1 (2012), 1. https://doi.org/10.1186/1297-9716-43-1

[28] Nicola De Maio, Colin J. Worby, Daniel J. Wilson, and Nicole Stoesser. 2018. Bayesian reconstruction of transmission within outbreaks using genomic variants. PLoS Comput Biol 14, 4 (2018), e1006117.

[29] Scott McGinnis and Thomas L Madden. 2004. BLAST: at the core of a powerful and diverse set of sequence analysis tools. Nucleic acids research 32, Web Server issue (07 2004), W20–W25. https://doi.org/10.1093/nar/gkh435

[30] Andrew Melnyk, Sergey Knyazev, Fredrik Vannberg, Leonid Bunimovich, Pavel Skums, and Alex Zelikovsky. 2020. Using earth mover’s distance for viral outbreak investigations. BMC Genomics 21, 5 (2020), 582. https://doi.org/10.1186/s12864-020-06982-4

[31] Hesam Montazeri, Susan Little, Niko Beerenwinkel, and Victor DeGruttola. 2018. Bayesian reconstruction of HIV transmission trees from viral sequences and uncertain infection times. arXiv:1801.07660 (2018).

[32] Marco J. Morelli, Gaёl Thébaud, Joёl Chadœuf, Donald P. King, Daniel T. Haydon, and Samuel Soubeyrand. 2012. A Bayesian Inference Framework to Reconstruct Transmission Trees Using Epidemiological and Genetic Data. PLoS Comput Biol 8, 11 (2012), e1002768.

[33] M. Otero and H.G. Solari. 2010. Stochastic eco-epidemiological model of dengue disease transmission by Aedes aegypti mosquito. Mathematical Biosciences 223, 1 (2010), 32–46.

[34] Y. Rubner, C. Tomasi, and L. J. Guibas. 1998. A metric for distributions with applications to image databases, In Sixth International Conference on Computer Vision (IEEE Cat. No.98CH36271). Sixth International Conference on Computer Vision (IEEE Cat. No.98CH36271), 59–66. https://doi.org/10.1109/ICCV.1998.710701

[35] Yossi Rubner, Carlo Tomasi, and Leonidas J. Guibas. 2000. The Earth Mover’s Distance as a Metric for Image Retrieval. International Journal of Computer Vision 40, 2 (2000), 99–121. https://doi.org/10.1023/A:1026543900054

[36] Marcel Salathé, Maria Kazandjieva, Jung Woo Lee, Philip Levis, Marcus W. Feldman, and James H. Jones. 2010. A high-resolution human contact network for infectious disease transmission. PNAS 107, 51 (2010), 22020–22025.

[37] Guillaume Salha, Stratis Limnios, Romain Hennequin, Viet-Anh Tran, and Michalis Vazirgiannis. 2019. Gravity-Inspired Graph Autoencoders for Directed Link Prediction. Proceedings of the 28th ACM International Conference on Information and Knowledge Management (2019), 589–598.

[38] S. Salipante and B. Hall. 2011. Inadequacies of Minimum Spanning Trees in Molecular Epidemiology. Journal of Clinical Microbiology 49 (2011), 3568–3575.

[39] Yuelong Shu and John McCauley. 2017. GISAID: Global initiative on sharing all influenza data - from vision to reality. Euro surveillance: bulletin Europeen sur les maladies transmissibles = European communicable disease bulletin 22, 13 (03 2017), 30494. https://doi.org/10.2807/1560-7917.ES.2017.22.13.30494

[40] Pavel Skums, Alex Zelikovsky, Rahul Singh, Walker Gussler, Zoya Dimitrova, Sergey Knyazev, Igor Mandric, Sumathi Ramachandran, David Campo, Deeptanshu Jha, Leonid Bunimovich, Elizabeth Costenbader, Connie Sexton, Siobhan O’Connor, Guo-Liang Xia, and Yury Khudyakov. 2018. QUENTIN: reconstruction of disease transmissions from viral quasispecies genomic data. Bioinformatics 34, 1 (5/15/2021 2018), 163–170. https://doi.org/10.1093/bioinformatics/btx402

[41] Enea Spada, Luciano Sagliocca, John Sourdis, Anna Rosa Garbuglia, Vincenzo Poggi, Carmela De Fusco, and Alfonso Mele. 2004. Use of the minimum spanning tree model for molecular epidemiological investigation of a nosocomial outbreak of hepatitis C virus infection. Journal of clinical microbiology 42, 9 (09 2004), 4230–4236. https://doi.org/10.1128/JCM.42.9.4230-4236.2004

[42] S Wohl, HC Metsky, SF Schaffner, A Piantadosi, M Burns, JA Lewnard, and et al. 2020. Combining genomics and epidemiology to track mumps virus transmission in the United States. PLoS Biol 18, 2 (2020), e3000611.

[43] Colin J. Worby, Marc Lipsitch, and William P. Hanage. 2014. Within-Host Bacterial Diversity Hinders Accurate Reconstruction of Transmission Networks from Genomic Distance Data. PLoS Comput Biol 10, 3 (2014), e1003549.

[44] Colin J Worby, Marc Lipsitch, and William P Hanage. 2017. Shared Genomic Variants: Identification of Transmission Routes Using Pathogen Deep-Sequence Data. American Journal of Epidemiology 186, 10 (5/15/2021 2017), 1209–1216. https://doi.org/10.1093/aje/kwx182

[45] Colin J Worby, Philip D O’Neill, Theodore Kypraios, Julie V Robotham, Daniela De Angelis, Edward J P Cartwright, Sharon J Peacock, and Ben S Cooper. 2016. Reconstructing transmission trees for communicable diseases using densely sampled genetic data. The annals of applied statistics 10, 1 (03 2016), 395–417. https://doi.org/10.1214/15-aoas898

[46] Xiao-Ke Xu, Xiao Fan Liu, Ye Wu, Sheikh Taslim Ali, Zhanwei Du, Paolo Bosetti, Eric H Y Lau, Benjamin J Cowling, and Lin Wang. 2020. Reconstruction of Transmission Pairs for Novel Coronavirus Disease 2019 (COVID-19) in Mainland China: Estimation of Superspreading Events, Serial Interval, and Hazard of Infection. Clinical Infectious Diseases 71, 12 (5/15/2021 2020), 3163–3167. https://doi.org/10.1093/cid/ciaa790

[47] R. J. F. Ypma, A. M. A. Bataille, A. Stegeman, G. Koch, J. Wallinga, and W. M. van Ballegooijen. 2012. Unravelling transmission trees of infectious diseases by combining genetic and epidemiological data. Proc. R. Soc. B. 279, 1728 (2012), 444–450.

[48] Rolf J F Ypma, W Marijn van Ballegooijen, and Jacco Wallinga. 2013. Relating phylogenetic trees to transmission trees of infectious disease outbreaks. Genetics 195, 3 (2013), 1055–62.

[49] Xu-Sheng Zhang and Daniela De Angelis. 2016. Construction of the influenza A virus transmission tree in a college-based population: co-transmission and interactions between influenza A viruses. BMC Infectious Diseases 16, 1 (2016), 38. https://doi.org/10.1186/s12879-016-1373-x

